# Impaired store-operated Ca^2+^ entry in mouse epidermis leads to reduced epidermal barrier function via Klk activation

**DOI:** 10.1101/2025.10.14.681588

**Authors:** Kaho Mochimatsu, Naoto Haruyama, Zhongze Liang, Masamitsu Nakanishi, Fumie Terao, Kanako Miyazaki, Keigo Yoshizaki, Gaku Tsuji, Takeshi Nakahara, Ichiro Takahashi

**Author notes:** Corresponding author Naoto Haruyama, DDS, Ph.D.

## Abstract

Store-operated Ca^2+^ entry (SOCE) is a mechanism by which STIM1/2 detect calcium depletion in the endoplasmic reticulum (ER) and activate ORAI1/TRPC channels on the cell membrane to induce calcium influx into the cytoplasm. This study aimed to investigate SOCE’s role in the skin epidermis *in vivo*, using female mice with epithelial tissue-specific Stim1 and Stim2 knockout (*Stim1/2 cKO*) and control mice (*Stim1/2^fl/fl^*). Keratinocytes from *Stim1/2 cKO* mice exhibited reduced Stim1 and Stim2 gene expression levels and impaired SOCE function compared with the controls. Histological analysis revealed hyperkeratosis in the *Stim1/2 cKO* tissues; however, no significant changes were observed in the proliferation or migration of keratinocytes or wound healing of the back skin. RNA-seq analysis indicated altered keratinization and cell-to-cell adhesion in the *Stim1/2 cKO* mice. Additionally, transepidermal water loss (TEWL) was increased significantly, and a biotinylated reagent diffused from the subcutaneous area into the granular layer, signifying compromised barrier function in the *Stim1/2 cKO* mice. The *Stim1/2 cKO* mice exhibited alterations in desmoglein 1 (Dsg1) and elevated levels of Kallikrein-related peptidase (Klk) 6 and Klk7, leading to increased trypsin- and chymotrypsin-like serine protease activities, respectively. These results suggest that SOCE dysfunction leads to hyperkeratosis and impaired epidermal barrier function via Klk activation without affecting skin cell proliferation *in vivo*. These novel findings improve understanding of the molecular mechanisms underlying calcium signaling in the epidermis barrier and suggest potential avenues for investigating related skin pathologies linked to calcium homeostasis, such as Hailey–Hailey disease (HHD) and Darier disease (DD).

## Introduction

Calcium ions (Ca^2+^) serve critical roles in epidermal proliferation, differentiation, and barrier functions *in vitro* and *in vivo*^1,2^. In the basal layer of the skin epidermis, the Ca^2+^ concentration is kept low and increases towards the upper layers, such as the stratum spinosum, stratum granulosum, and stratum corneum. This Ca^2+^ concentration gradient is thought to be essential for normal epidermal growth and differentiation^3,4^. However, recent studies show that the majority of free Ca^2+^ measured in the epidermis is derived from intracellular Ca^2+^ stores such as the endoplasmic reticulum (ER) and Golgi apparatus^5^. Therefore, the fine regulation of intracellular Ca^2+^ localization could be important to maintaining epidermal homeostasis and differentiation.

Hailey–Hailey disease (HHD) stems from mutations in the *ATP2C1* gene, encoding the ATP pump SPCA1 responsible for sequestering calcium in the Golgi apparatus^6^.

Disruptions of the intracellular Ca²⁺ store caused by mutations in the *ATP2C1* gene affect the assembly and trafficking of desmosomal proteins, such as desmoplakin and desmoglein 3, to the cell membrane in HHD^7,8^. In contrast, Darier disease (DD) results from mutations in the *ATP2A2* gene, encoding the calcium pump SERCA2 in the ER ^9,10^. Defective ER calcium homeostasis can disrupt cell-to-cell adhesion through the improper reorganization of junctional components^9,11,12^. HHD keratinocytes exhibit a diminished response to heightened extracellular Ca²⁺, leading to decreased intracellular Ca²⁺ relative to controls^6^. Conversely, SERCA2 dysfunction in DD patients causes a reduction in ER Ca²⁺ content and increased cytosolic Ca²⁺ ^13^. Although HHD and DD display distinct intracellular Ca²⁺ levels and responses to extracellular Ca²⁺, desmosomal adhesion may be a shared challenge, suggesting that finely regulated intracellular Ca²⁺ localization is essential for keratinocyte adhesion and epidermal integrity.

Store-operated Ca^2+^ entry (SOCE) is a major mechanism by which calcium enters non-excitable and excitable cells^14,15^. SOCE is a phenomenon in which the ER membrane molecules STIM1 and STIM2, upon sensing a decrease in the calcium concentration in the ER, translocate to the plasma membrane and form channels with TRPC and/or ORAI1 molecules to generate a highly calcium-selective current (ICRAC) and induce calcium influx^16,17^. A recent study on *Stim1* knockout mice has shown that impaired Stim1 results in accelerated wound healing based on disruption of the epidermal barrier via tape stripping^18^. On the other hand, epidermal atrophy and sporadic alopecia have been reported in *Oral1* knockout mice^19^. Therefore, SOCE’s function in epidermal skin appears to be inconsistent and requires further elucidation, especially *in vivo*.

In this study, impaired SOCE function and altered intracellular Ca^2+^ concentration were hypothesized to potentially cause changes in epithelial cell proliferation and the epidermal skin barrier. Here, the skin tissue was analyzed from ablated *Stim*1 and *Stim*2 genes by crossing the *Stim1* and/or *Stim2* floxed mice with *Keratin14 (K14)-Cre* transgenic mice^20^. The epidermal barrier function was impaired, possibly resulting from the disruption of desmoglein 1 (Dsg1) via Kallikrein-related peptidase (Klk) activation.

## Results

### Conditional ablation of *Stim1/2* in the mouse epidermis

First, the Cre-mediated site-specific recombination was assessed using β-gal staining in *K14-Cre* mice, since the *K14-Cre* mice harbored the ROSA-stop-LacZ cassette. The epidermis and hair follicles showed strong β-gal activity, confirming successful Cre recombination in the skin (Fig. 1a). In the following experiments, *Stim1/2^fl/fl^*mice without expressed *K14-Cre* were used as controls *(Stim1/2^fl/fl^*), and *Stim1/2^fl/fl^ K14-Cre* mice were used as conditional knockout mice (*Stim1/2 cKO*). Immunohistochemistry revealed that the Stim1 protein was present in all layers of the epidermis and hair follicles in the control *Stim1/2^fl/fl^* mice, whereas fluorescence intensity was markedly reduced in *Stim1/2 cKO* mice, confirming the targeted ablation (Fig. 1b). The quantitative PCR (qPCR) analysis showed a significant decrease in the expression levels of Stim1 and Stim2 mRNA in the *Stim1/2 cKO* group compared with the controls (Fig. 1c). Correspondingly, the protein expression levels of Stim1 and Stim2 were diminished in *Stim1/2 cKO* mice (Fig. 1d and Supplemental Material), confirming the effective knockout of the *Stim1* and *Stim2* genes.

**Figure 1.**
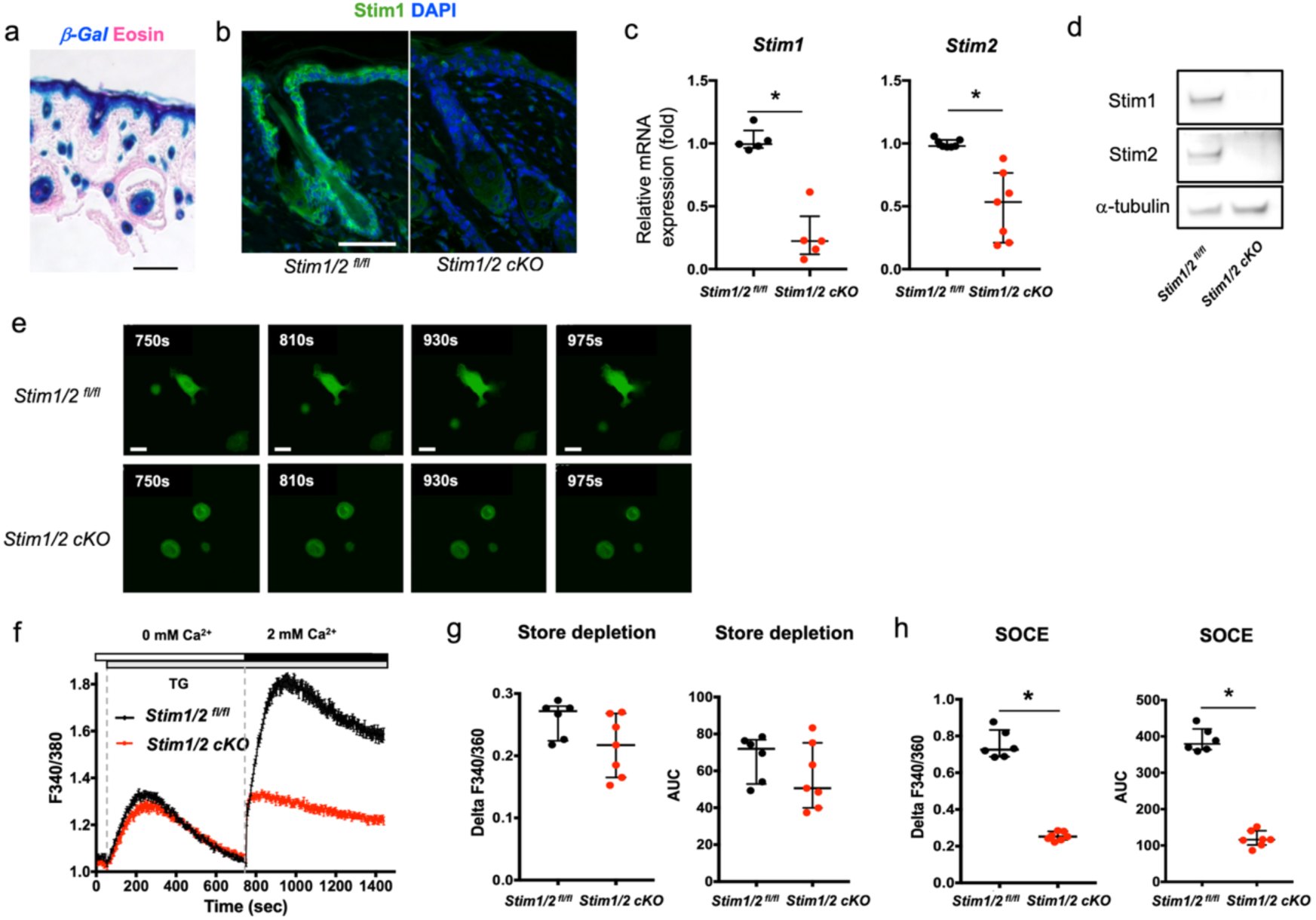
Conditional ablation of *Stim1/2* in the mouse epidermis leads to impaired function of SOCE. **(a)** A skin tissue section with β-gal staining of *K14-Cre* mice. *LacZ* activity was found in the epidermis and hair follicles. The section was counterstained with eosin. Bar = 50 μm. **(b)** Immunofluorescence staining of Stim1 (green) in back skin sections from *Stim1/2^fl/fl^* control (left panel) and *Stim1/2 cKO* (right panel) 6-week-old littermate mice. Nuclei were counterstained with DAPI (blue). The localization of Stim1 was found in most layers of the epidermis, except the stratum corneum, and hair follicles in *Stim1/2^fl/fl^*mice. However, the fluorescence intensity from Stim1 decreased dramatically in the *Stim1/2 cKO* epidermis. Bars = 50 μm. **(c)** qPCR from 6-week-old mouse skin. Fold change in gene expression was calculated using the ΔΔCT method, normalizing to *Hprt* and then the *Stim1/2^fl/fl^* control mice (data represent median with interquartile range, n = 5–6/genotype). The gene expression levels of *Stim1* and *Stim2* were significantly decreased in the *Stim1/2 cKO* epidermis. *p-*value: calculated using the Mann–Whitney U test; *: statistically significant (p < 0.05). **(d)** Western blotting from 6-week-old littermate mice skin. The panel shows representative data collected from four independent pairs of littermate mice. α-tubulin was loaded as an internal control. **(e)** Fluorescence imaging of intracellular Ca^2+^ levels by Fluo4. After depleting calcium from the ER by adding 2 μM TG, 2 mM CaCl_2_ was added to the keratinocytes (750 seconds after TG stimulation), and the fluorescence was monitored using time-lapse microscopy. The signal intensity increased in the *Stim1/2^fl/fl^* keratinocytes (upper panels) until 975 seconds. However, minimal changes in signal intensity were found in the cells from *Stim1/2 cKO* mice (lower panels). Bars = 20 μm. **(f)** Quantification of intracellular Ca^2+^ levels by Fura2. Upon stimulation with TG (at 50 seconds), both *Stim1/2^fl/fl^* (black tracing) and *Stim1/2 cKO* keratinocytes (red tracing) showed significant, though comparable, releases of Ca^2+^ from the ER. In contrast, stimulation with 2 mM Ca^2+^ (at 750 seconds) induced significant Ca^2+^ entry in *Stim1/2^fl/fl^*(black tracing) relative to *Stim1/2 cKO* keratinocytes (red tracings). **(g-h)** Statistical analysis of the average delta peak and the AUC for TG-induced store depletion (g) and for SOCE (h), respectively, as shown in (f). Significant differences were found in Ca^2+^ influx (SOCE), but not in Ca^2+^ release from the ER (store depletion). (Data represent median with interquartile range, n = 7/genotype). *p-*value: calculated using the Mann–Whitney U test; *: statistically significant (p < 0.05).

### Knockout of *Stim1/2* impaired the function of SOCE in the epidermis

Calcium imaging and intracellular Ca^2+^ level quantification were performed using Fluo4 and Fura2, respectively (Fig. 1e and f). After depleting Ca^2+^ in the ER via thapsigargin (TG), the intracellular Ca^2+^ levels in the *Stim1/2^fl/fl^*-derived mouse keratinocytes appeared to increase in response to adding 2 mM extracellular calcium (Fig. 1e, upper panels) compared with the cells from *Stim1/2 cKO* mice (Fig. 1e, lower panels).

Quantitative analysis using Fura2 indicated that TG-induced ER calcium depletion led to a modest increase in intracellular Ca^2+^ for both genotypes, followed by a decrease (Fig. 1f, 50–750 seconds). However, a significant Ca^2+^ influx was observed in *Stim1/2^fl/fl^* keratinocytes after adding 2 mM calcium, while the response in *Stim1/2 cKO* keratinocytes was markedly diminished (Fig. 1f, 750 seconds∼). These results indicate that SOCE function is significantly impaired in the epidermis of *Stim1/2 cKO* mice.

### *Stim1/Stim2 cKO* results in hyperkeratosis in the back skin

Macroscopically, no significant differences in whiskers or dorsal coat hair were observed between the genotypes (Fig. 2a, upper panels). However, the ventral skin exhibited a red coloration, accompanied by reduced hair density in the *Stim1/2 cKO* mice (Fig. 2a, lower panels). Moreover, desquamation was evident when the dorsal coat hair was clipped in the *Stim1/2 cKO* mice (Fig. 2b, arrows). Histological analysis using hematoxylin and eosin (H&E) staining revealed epidermal thickening in *Stim1/2 cKO* mice compared with *Stim1/2^fl/fl^*mice (Fig. 2c). Examination of E-cadherin (Cdh1) and filaggrin (Flg) expression in skin sections showed a significant increase in the thickness of both Cdh1- and Flg-positive cell layers in the *Stim1/2 cKO* mice. Notably, Flg-positive layers, indicative of the stratum granulosum and stratum corneum, increased dramatically (Fig. 2d and e). These results suggest that the *Stim1/2 cKO* mice had thickened skin, resulting in hyperkeratosis.

**Figure 2.**
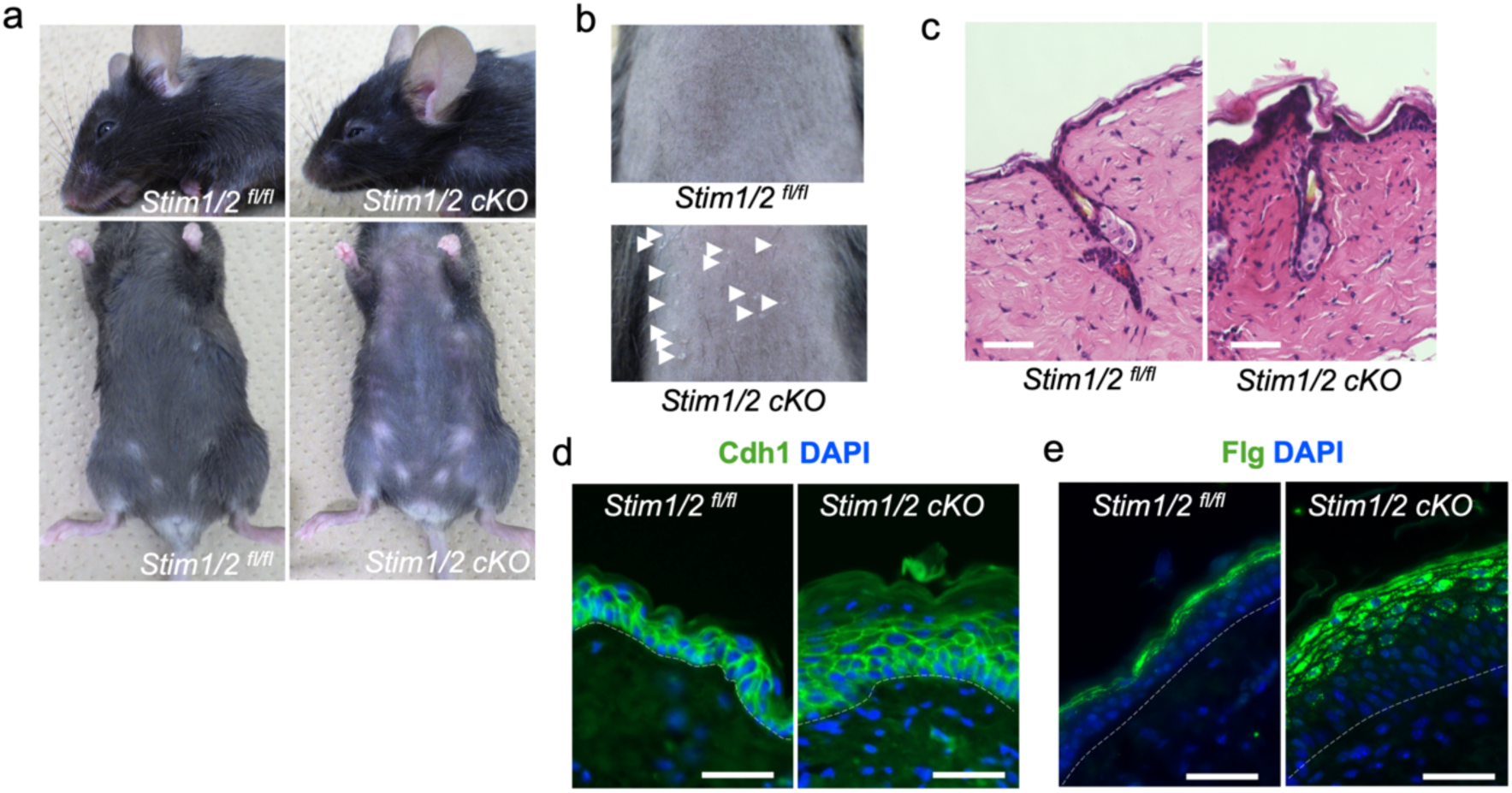
*Stim1/Stim2 cKO* results in hyperkeratosis in the back skin. **(a-b)** Macroscopic observation of 6-week-old control and *Stim1/2 cKO* mice. (a) No significant differences in whisker or dorsal coat hair between genotypes (upper panels); the ventral skin exhibited red coloration, accompanied by a low hair density in the *Stim1/2 cKO* mice (lower panels). (b) Desquamation was evident on the dorsal skin (arrows). The back skin was shaved before imaging with electric clippers. The changes in (a) and (b) were characteristic of all 6-week-old *Stim1/2 cKO* mice analyzed in this study. **(c)** H&E staining of back skin paraffin sections from control mice (left panel) and *Stim1/2 cKO* littermate mice (right panel). Epidermal thickness and keratinization increased in *Stim1/2 cKO* mice. Bars = 50 μm. **(d-e)** Immunofluorescence staining of Cdh1 (green in panel d) and Flg (green in panel e) in back skin sections from control (left panels) and *Stim1/2 cKO* littermate mice (right panels). The thickness of both the Cdh1- and Flg-positive cell layers increased in the *Stim1/2 cKO* mice. Notably, the Flg-positive cell layers, corresponding to the stratum granulosum and stratum corneum, increased dramatically (e). Nuclei are counterstained with DAPI (blue). Bars = 30 μm.

### *Stim1/Stim2 cKO* mice show no changes in cell proliferation or migration in keratinocytes or the back skin wound healing

To elucidate the cause of the observed epidermal thickening in *Stim1/2 cKO* mice, tissue sections for the nuclear protein Ki67 were examined, a marker of cell proliferation, using immunohistochemistry (IHC). The number of Ki67-positive cells did not differ between the genotypes (Fig. 3a and b), and *in vitro* proliferation assays confirmed no significant differences over four days (Fig. 3c). A scratch assay indicated comparable migratory abilities between genotypes (Fig. 3d and e). Additionally, full-thickness excisional wounds were created on the dorsal skin, and wound closure was monitored over eight days in both genotypes. No significant differences in wound healing speed were observed (Fig. 3f and g), suggesting that the deletion of *Stim1/2* does not impair keratinocyte proliferation or migration *in vivo* or *in vitro*.

**Figure 3.**
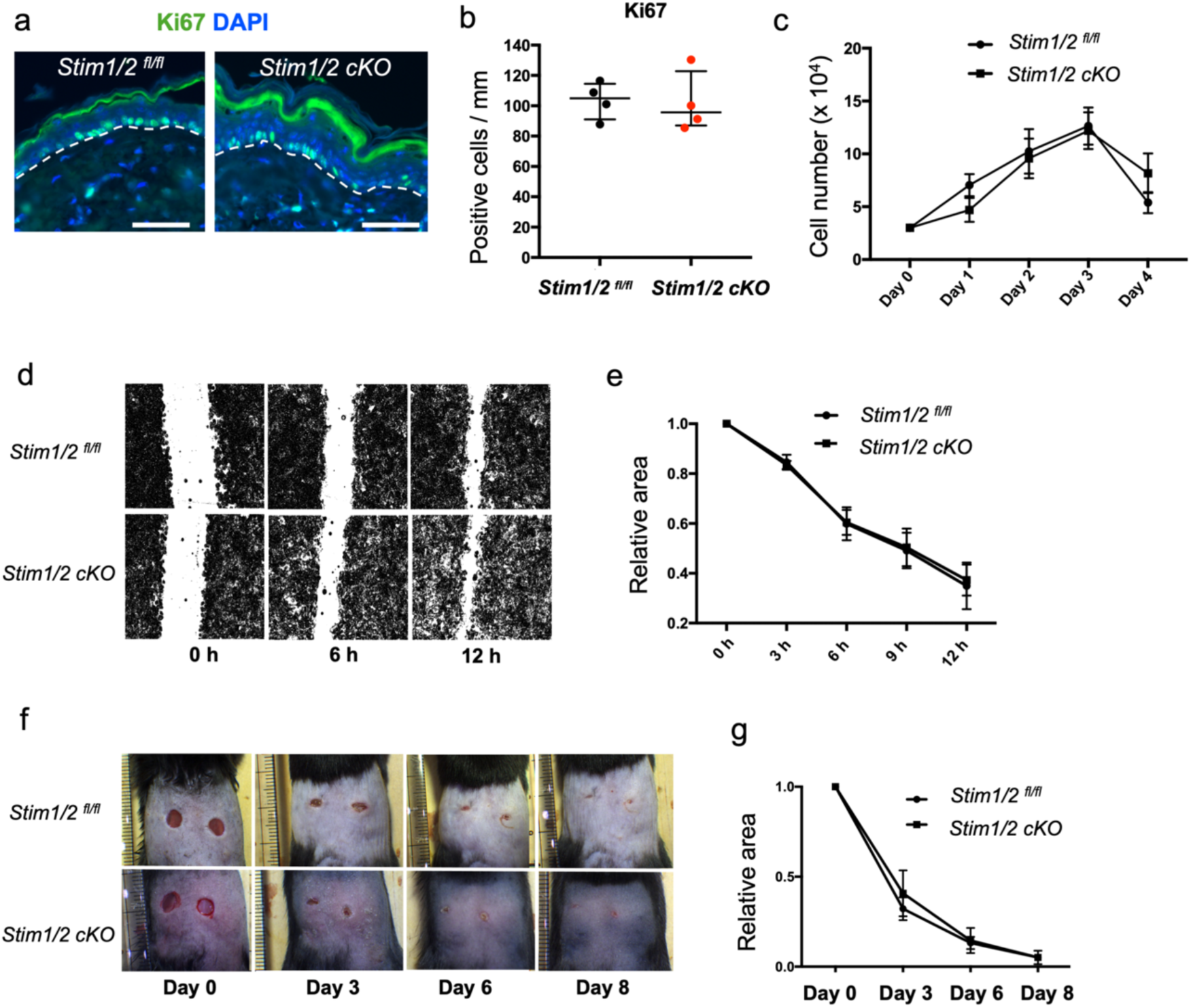
*Stim1/Stim2 cKO* mice show no changes in cell proliferation and migration in keratinocytes or the back skin wound healing. **(a)** Immunofluorescence staining of Ki67 (green) in back skin sections from 6-week-old *Stim1/2^fl/fl^* control (left panel) and *Stim1/2* cKO (right panel) littermate mice. Nuclei are counterstained with DAPI (blue). Ki67 localization was observed in the nuclei of the basal layer (stratum basale) in the epidermis. The number of fluorescence-positive cells appeared to be equivalent in both genotypes. Strong nonspecific signals were observed in the keratinized layers (stratum corneum). Bars = 50 μm. **(b)** The graph shows the quantification of Ki67 staining of control skin (n = 4) and *Stim1/2 cKO* skin (n = 4) as the number of Ki67-positive cells/mm of basal layer. (Data represent median with interquartile range, n = 4/genotype). *p-*value: calculated using the Mann–Whitney U test. **(c)** Cell proliferation curves of mouse keratinocytes *in vitro*. No significant differences were found in the cell numbers between the *Stim1/2^fl/fl^* (round) and *Stim1/2 cKO* mice (square) derived cells from Day 0 to 4. Cell numbers were counted directly over four days **(d)** using an *in vitro* wound healing assay to assess the cell migration of keratinocytes. Representative black-and-white images were converted from bright-field images using ImageJ. Gaps (white areas) created by straight-line scratches across the keratinocyte monolayer (black areas) were monitored for 12 hours. The gap distances showed similar migratory abilities in cells isolated from *Stim1/2^fl/fl^* (upper panels) and *Stim1/2cKO* (lower panels) mice. **(e)** Quantitative analysis of the gap area shown in (d). No significant differences were found between the *Stim1/2^fl/fl^* (round) and *Stim1/2* cKO (square) derived cells. **(f)** *In vivo* wound healing of the mouse back skin and **(g)** the quantitative analysis of the wound area. No significant differences were found in the wound recovery (area) between the *Stim1/2^fl/fl^* (round) and *Stim1/2 cKO* (square) back skin over eight days. **(c, e,** and **g)** Data represent mean ± SD (n = 5–6/genotype).

### Expression profile analysis by RNA-seq suggested the altered keratinization and cell-to-cell adhesion

The gene expression profile was assessed using RNA-seq to understand the phenotypic changes observed. A total of 21 upregulated and 26 downregulated genes were identified in the *Stim1/2 cKO* mice (Fig. 4a). Gene ontology (GO) analysis revealed that several critical skin biology processes were altered by impaired SOCE in keratinocytes (Fig. 4b and Supplemental Table 2). These processes include intermediate filament organization (GO:0045109), supramolecular fiber organization (GO:0097435), protein autoprocessing (GO:0016540), and epidermis development (GO:0008544). Keratins, the major intermediate filaments, form a dense mesh of cytoplasmic fibers linked to desmosomes at cell-to-cell junctions and to hemidesmosomes at the cell–substrate interface ^21^. Therefore, it was hypothesized that modulated intermediate filament functions and/or desmosomal junctions would affect the epidermal barrier function because of *Stim1/2* deletion.

**Figure 4.**
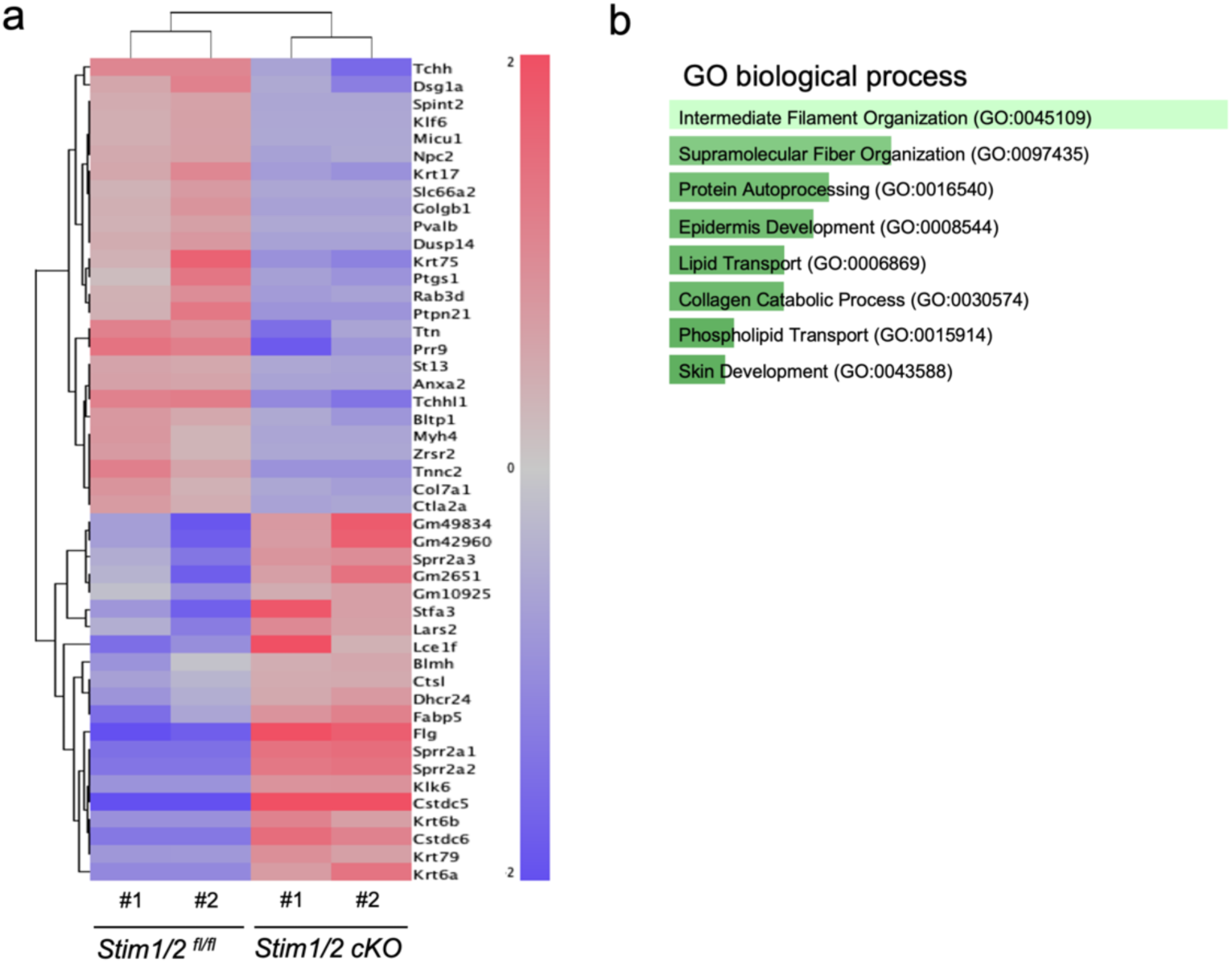
Gene expression analysis by RNA-seq suggested altered keratinization and skin development. **(a)** Gene expression profiles by RNA-seq. The heatmap showing z-scores of genes differentially expressed by two-fold, adjusted *p* < 0.05, identified 21 upregulated and 26 downregulated genes in the *Stim1/2 cKO* mice. **(b)** GO biological process terms. The results include intermediate filament organization (GO:0045109), supramolecular fiber organization (GO:0097435), protein autoprocessing (GO:0016540), and epidermis development (GO:0008544).

### *Stim1/Stim2 cKO* mice show reduced barrier function in the epidermis

In the course of monitoring the older mice (77-week-old), notable desquamation of the dorsal skin, sporadic alopecia from the ocular region to the dorsal coat, and dry skin on the flank were observed in the *Stim1/2 cKO* mice (Fig. 5a). In younger mice (6-week-old), treated with acetone to challenge the skin barrier, the *Stim1/2 cKO* mice exhibited substantial keratinization compared with the control mice (Fig. 5b, arrows), indicating an altered epidermal response to barrier disruption. Furthermore, transepidermal water loss (TEWL), an indicator of skin barrier integrity, was significantly higher in the *Stim1/2* cKO mice (Fig. 5c), indicating impaired barrier function. To assess the skin barrier status, an *in vivo* diffusion assay of Lucifer yellow and biotin was performed to examine the outside-in and inside-out barrier function, respectively ^22^. Lucifer yellow penetration via the stratum corneum barrier was not observed in either *Stim1/2^fl/fl^* or *Stim1/2 cKO* mice (Fig. 5d). However, in the *Stim1/2 cKO* mice, the biotinylation reagent diffused from the subcutaneous area through the epidermis to the outermost layer containing nuclei (the stratum granulosum) (Fig. 5e, arrows). Notably, the biotinylation reagent did not display apparent diffusion along with the cell membrane of the stratum granulosum in the *Stim1/2^fl/fl^*mice compared with the *Stim1/2 cKO* mice. This diffusion above the stratum granulosum suggests a disrupted tight junction barrier^23^. These findings demonstrate that the absence of Stim1 and Stim2 in keratinocytes compromises inside-out skin barrier function *in vivo*.

**Figure 5.**
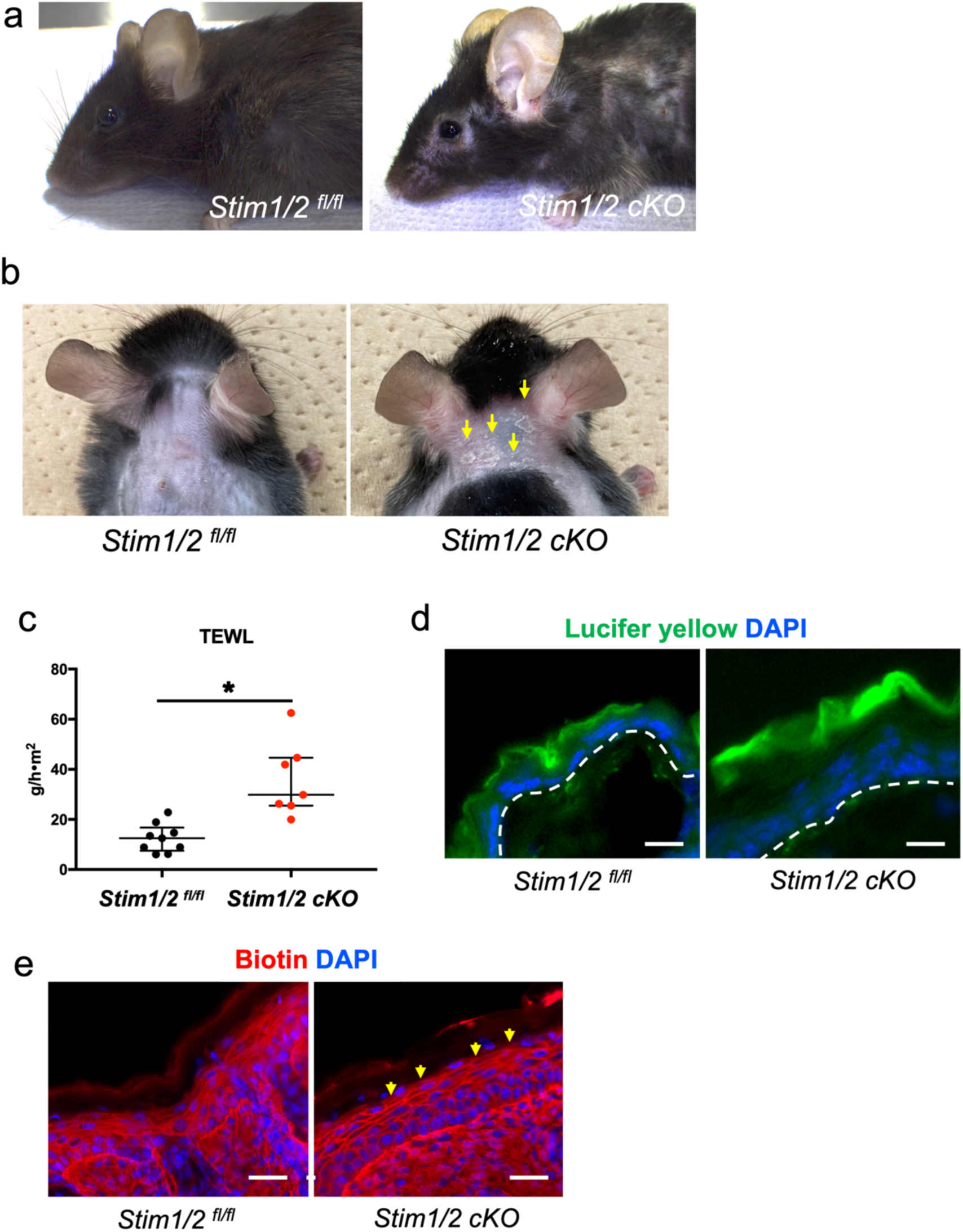
*Stim1/Stim2 cKO* mice show reduced barrier function in the epidermis. **(a)** Macroscopic observation of 77-week-old *Stim1/2 ^fl/fl^* control and *Stim1/2 cKO* mice. Desquamation of the dorsal skin and sporadic alopecia were notable in the *Stim1/2 cKO* mice. **(b)** Acetone-induced skin barrier disruption. The day after treatment with acetone-soaked cotton balls, a substantial increase in keratinization was observed in *Stim1/2 cKO* mice compared with *Stim1/2^fl/fl^*mice (see arrows). **(c)** TEWL measurements performed on the lower back skin of *Stim1/2 ^fl/fl^* mice (black) and *Stim1/2 cKO* mice (red). Each dot represents the mean of three measurements for an individual mouse. The amount of TEWL was significantly increased in the *Stim1/2 cKO* mice. These data represent median values with interquartile ranges (n = 7–9/genotype). *p-*value: calculated using the Mann–Whitney U test; *: statistically significant (*p* < 0.05). **(d)** *In vivo* diffusion of Lucifer yellow for inward barrier function. The fluorescent micrographs show the distribution of Lucifer yellow in the back skin of *Stim1/2 ^fl/fl^* and *Stim1/2 cKO* mice at 6 weeks of age. The Lucifer yellow reagent only diffused into a portion of the layers in the stratum corneum (cornified layer) but did not reach the stratum granulosum. Nuclei are counterstained with DAPI. Dashed line, basement membrane. Bars = 30 μm **(e)** *in vivo* diffusion assay of biotin for inside-out barrier function. The biotinylating reagent was injected subcutaneously to determine whether the diffusion of materials through the paracellular pathway was affected. An isotonic solution containing a biotinylating reagent was injected into the dermis on the backs of newborns. After 30 minutes of incubation, the skin was dissected and frozen. The sections were stained with Texas Red-conjugated streptavidin. In the *Stim1/2 cKO* mice, the biotinylating reagent diffused through the paracellular spaces from the stratum basale to the top layer of the stratum granulosum (arrows); however, this diffusion was not significant in the stratum granulosum as observed in the *Stim1/2 ^fl/fl^* mice. Nuclei are counterstained with DAPI (blue). Bars = 30 μm.

### *Stim1/2* cKO mice display defective cell–cell adhesion marker desmogrein1 (Dsg1) and increased activities of the trypsin- and chymotrypsin-like proteases

Expression of the epidermal barrier marker Dsg1 was quantified to assess the epidermal barrier. RNA-seq analysis indicated that *Dsg1* levels were lower in the *Stim1/2 cKO* mice (Fig. 3a). The qPCR analysis further confirmed a decrease in *Dsg1* expression in five of six littermate pairs; however, the difference was not statistically significant (p = 0.14, Fig. 6a). Indeed, in the *Stim1/Stim2* cKO mice, Dsg1 showed a dramatically reduced fluorescence intensity, but no distinct difference in the expression pattern in the skin (Fig. 6b), suggesting that the protein related to cell-to-cell adhesion, Dsg1, is attenuated in *Stim1/Stim2 cKO* mice, possibly via transcriptional and post-translational processes.

**Figure 6.**
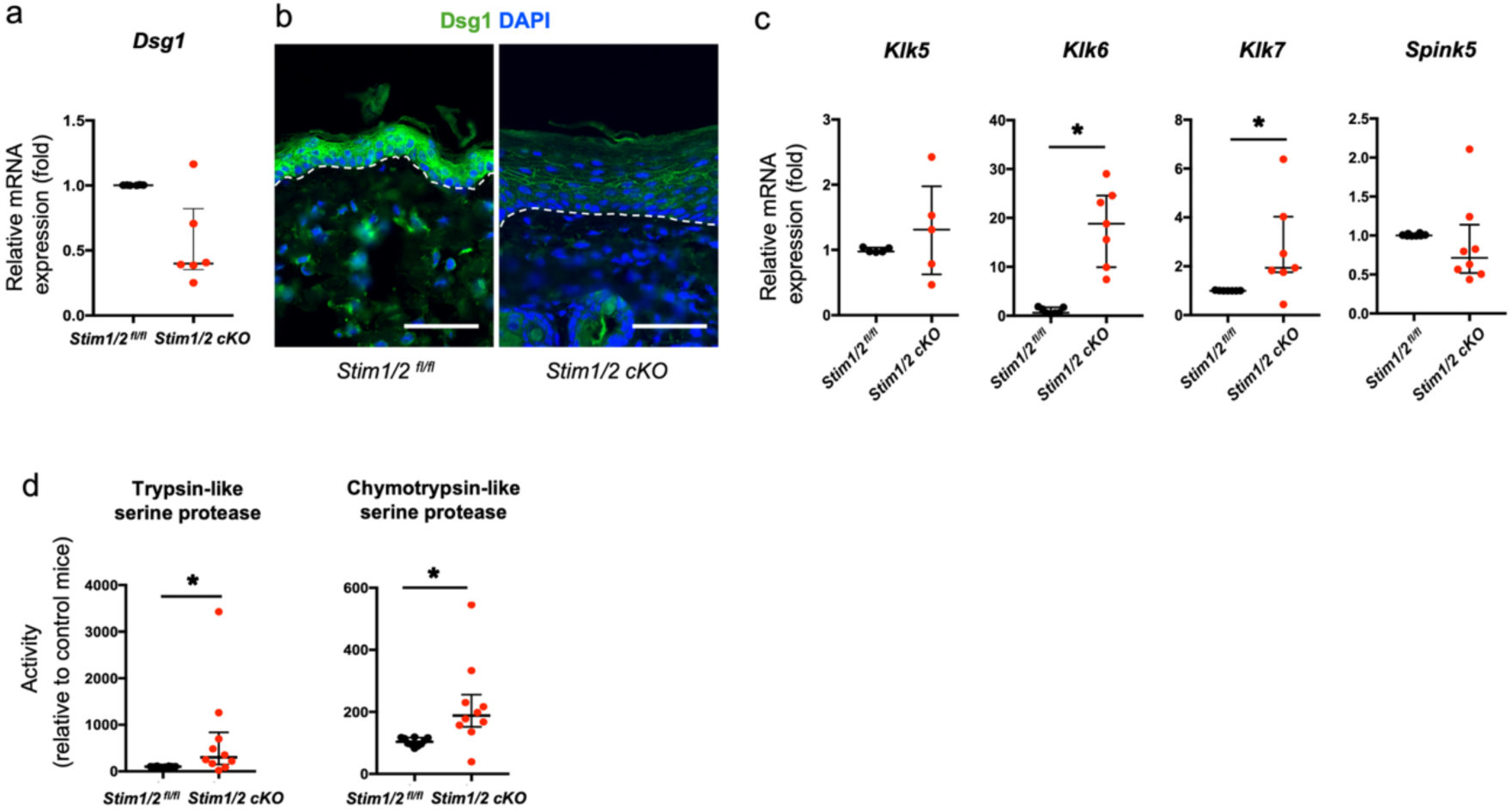
*Stim1/2 cKO* mice display defective cell–cell adhesion marker Dsg1 and increased activity of the trypsin- and chymotrypsin-like proteases in the skin tissue. **(a)** qPCR of Dsg1 in the skin of 6-week-old mice. The fold changes in gene expression levels were calculated using the 1¢1¢CT method, normalizing to *Hprt* and then the *Stim1/2^fl/fl^* control littermate mice (data represent median values with interquartile ranges, n = 6/genotype). Gene expression levels of *Dsg1* decreased in *Stim1/2 cKO* epidermis compared with *Stim1/2^fl/fl^* controls in five out of six littermates. However, the difference was not statistically significant. *p-*value: calculated using the Mann–Whitney U test. **(b)** Immunofluorescence staining of desmogrein1 (Dsg1) (green) in back skin sections from 6-week-old *Stim1/2^fl/fl^* control (left panel) and *Stim1/2* cKO (right panel) littermate mice. Nuclei are counterstained with DAPI (blue). Dsg1 localization was found in all layers of the epidermis and hair follicles in *Stim1/2^fl/fl^* mice; however, the fluorescence intensity from Stim1 decreased dramatically in the *Stim1/2 cKO* epidermis. Bars = 50 μm. **(c)** qPCR of *Klk5*, *Klk6*, *Klk7*, and *Spink5* in the skin of 6-week-old mice. The fold changes in gene expression levels were calculated using the 1¢1¢CT method, normalizing to *Hprt* and then the *Stim1/2^fl/fl^* control littermate mice (data represent median values with interquartile ranges, n = 6–8/genotype). The expression levels of *Klk6 and Klk7* showed significant increases in the *Stim1/2 cKO* epidermis compared with the control. *p-*value: calculated using the Mann–Whitney U test. *: statistically significant (*p* < 0.05). **(d)** Activities of trypsin- and chymotrypsin-like serine proteases in the epidermis. Measurements of trypsin-like serine protease activity in epithelial protein extracts from the back skin of *Stim1/2^fl/fl^* controls (black) and *Stim1/2 cKO* (red) mice were performed using the fluorogenic substrate for trypsin-like serine proteases Boc-Phe-Ser-Arg-MCA (left) and the chymotrypsin-like serine proteases Suc-Leu-Leu-Val-Tyr-MC (right). The fluorescence intensities of the negative controls (proteins with PMSF) were subtracted from those of the samples (proteins without PMSF) and normalized to the *Stim1/2^fl/fl^*mice (data represent median values with interquartile ranges, n = 10/genotype). *p-*value: calculated using the Mann–Whitney U test. *: statistically significant (p < 0.05).

Since the fluorescence intensity of Dsg1 was reduced in the IHC of *Stim1/2 cKO* mice, it was hypothesized that Klk5, Klk6, and Klk7, the major serine proteases in skin ^24^, promote stratum corneum exfoliation and barrier disruption by degrading desmosomal cadherin in the epidermis. The expression levels of *Klk5, Klk6, Klk7,* and *Spink5*, encoding LEKTI1, an inhibitor of Klk activity in mouse epidermis, were confirmed using qPCR. The *Klk5* and *Spink5* mRNA expression levels did not differ significantly between groups. In contrast, the expression of *Klk6* and *Klk7* was increased in *Stim1/2 cKO* mice (Fig. 6c). Indeed, the increased expression of *Klk6* was indicated by the RNA-seq results (Fig. 3a). Trypsin- and chymotrypsin-like serine protease activities in the epidermis were further examined, representing the Klk5 or Klk6 and Klk7 activity, respectively. Interestingly, both activities were significantly increased in *Stim1/Stim2 cKO* mice compared with the control (Fig. 6d), suggesting that impaired intracellular Ca²⁺ influx (reduced SOCE) may alter trypsin- and chymotrypsin-like activities, potentially affecting Klk6 and Klk7. This alteration likely contributes to the degradation of Dsg1 and subsequent epidermal barrier disruption.

## Discussion

Ca^2+^ ions have been suggested to be strongly associated with epidermal proliferation, differentiation, and barrier homeostasis ^25,26^. A recent clinical study revealed that patients with genetically confirmed DD exhibited reduced STIM1 localization in their skin specimens, although this was not observed in HHD specimens^27^. In this study, impaired function of SOCE in the epidermis was clearly demonstrated to lead to hyperkeratosis and reduced barrier function in the mouse skin without altering proliferation. The localization of desmosomal cadherin Dsg1 was altered, potentially via an increase in serine protease activity in the *Stim1/2 cKO* mice. These results provide clear evidence indicating that SOCE regulated by Stim1/2 plays a major role in maintaining skin barrier homeostasis *in vivo*. Considering that it was hypothesized that the epithelial cell proliferation and barrier function would be impaired, this hypothesis was partially accepted.

Two STIM homologs exist in mammals: STIM1 and STIM2. STIM1 functions as a primary ER calcium sensor, activated by calcium release that subsequently opens ORAI1/TRPC channels in the plasma membrane. In contrast, the role of STIM2 on SOCE has been suggested to be modest ^28–32^, depending on the cell line type or experimental conditions. Therefore, in this mouse model, both Stim1 and Stim2 were knocked out, and calcium imaging of keratinocytes was performed to examine the extent of functional changes resulting from both *Stim1/2* gene ablation. The knockout of *Stim1/2* significantly, but not completely, suppressed Ca²⁺ influx, indicating their prominent role in SOCE in response to store depletion in keratinocytes (Fig. 1f).

Unexpectedly, the knockout of *Stim1/2* did not reduce TG-induced passive Ca²⁺ release from ER stores in low Ca²⁺ media, suggesting that the *Stim1/2* knockout does not substantially affect basal ER Ca²⁺ levels in these cells. These findings contradict previous reports using siRNA for *Stim1/2* in HaCaT cells ^33^ and findings on Stim1 single cKO mice^18^. To understand the cause of hyperkeratosis (epidermal thickening) in the *Stim1/2 cKO* mice, cell proliferation, migration, and wound healing were examined. Previous studies indicated that impaired SOCE leads to reduced keratinocyte proliferation and migration through Orai1 channel function *in vitro*^33,34^. However, no significant differences in these parameters or wound healing were observed between *Stim1/2^fl/fl^*and *Stim1/2 cKO in vivo*. These results contrast with previous reports on the *Stim1 cKO* mouse model^18^. The reasons for these discrepancies remain unclear, potentially linked to the differences in materials, experimental methods, or the gene of interest being knocked out.

The literature shows that changes in the concentrations of intracellular Ca²⁺ stores cause barrier disruption and abnormal keratinization, *e.g*., HHD, which typically appears during the second to fourth decades of life, and DD, which manifests in late adolescence or early adulthood ^35^. Reportedly, in normal human keratinocytes, inhibition of SERCA by TG impairs intracellular transport of desmoplakin, desmoglein, and desmocollin to the cell surface ^7^. Both HHD and DD demonstrate abnormal desmosomal adhesion ^7,12,36,37^. Additionally, the expressions of the tight junction proteins claudin-1 and claudin-4 were suggested to be regulated by the *ATP2C1* gene in cultured keratinocytes ^36^. Therefore, a change in intracellular Ca²⁺ resulting from altered SOCE would also alter cell-to-cell adhesion and the epithelial barrier, potentially through the transcription and post-translational processes of cell-to-cell junctional proteins in this study’s mouse model. Intracellular transport of Dsg1 under compromised SOCE will be tested using further *in vitro* experiments in the future.

KLK belongs to a family of serine proteases involved in cleaving desmosomal proteins either directly or by participating in proteolytic activation, and contributing to the epidermal desquamation^38,39^. Previous studies have shown that increased activities of the Dsg1-degrading enzymes, KLK5, KLK6, and KLK7, disrupt the skin’s barrier function ^39,40, 41^. The connection between changes in Ca^2+^ concentration and increased KLK activity remains poorly understood. A previous report suggested that increasing extracellular Ca^2+^ induced KLK5 and KLK7 expression *in vitro* ^42^. On the other hand, another *in vitro* report indicates that the activities of trypsin- and chymotrypsin-like serine proteases were significantly up- and downregulated by extracellular high Ca^2+^ levels, respectively, in a dose-dependent manner, in normal human epidermal keratinocytes^43^. In this study, Klk6 and Klk7 exhibited significantly increased gene expression levels. Moreover, an analysis of serine protease activity revealed increases in the activities of trypsin- and chymotrypsin-like serine proteases. Therefore, the hyperactivity of Klk6 and Klk7 appears to play a significant role in the development of barrier breakdown and desquamation in *Stim1/2 cKO* mice.

In conclusion, these results improve understanding of the molecular mechanisms underlying calcium signaling in the epidermis and suggest potential avenues for investigating related skin pathologies linked to calcium homeostasis, such as HHD and DD.

## Materials and methods

### Animal

Epithelium-specific *Stim1/2* knockout mice were used as reported previously^20^. Female mice (age, 6–8 weeks) were used in all experiments unless noted otherwise. All mice were housed in a specific pathogen-free facility and fed *ad libitum* a normal diet and sterilized water under a 12-hour light/dark cycle. All animal procedures and breeding programs were approved by the Kyushu University Animal Care and Use Committee (Protocol #A22-031, #A24-106) and complied with the ARRIVE guidelines.

### Histology, immunohistochemistry, Lac Z staining, Lucifer yellow assay, and biotin permeability assay

Back skin samples from mice were fixed in Bouin’s solution (Fujifilm Wako, Osaka, Japan), dehydrated in a graded ethanol series, embedded in paraffin, and sectioned into 5 µm slices for H&E staining or immunohistochemistry. Immunohistochemistry was performed using primary antibodies against Stim1, Dsg1, Cdh1, Flg, and Ki67.

Subsequently, the samples were prepared as follows: incubated with Alexa Fluor 488-conjugated secondary antibodies and counterstained with DAPI. Frozen skin sections embedded in OCT compound (Sakura Finetek, Tokyo, Japan) were obtained from 6- to 7-week-old, post-natal day 7, or neonatal mice for Lucifer yellow permeability assay, β-gal staining, or biotin permeability assay, respectively. See the Supplemental Methods section for details.

### Induction of barrier disruption by acetone and measurement of transepidermal water loss (TEWL)

The protocol for acetone-induced barrier disruption was performed as described previously^44^. The back skin hair was shaved, and the shaved upper back skin area was treated with acetone-soaked cotton balls for 5 minutes. One day later, the skins of the experimental animals were observed and recorded. TEWL was measured using a Vaposcan (AS-VT200RS, Asch Japan, Tokyo, Japan) on the lower back skin region without applying an acetone treatment. The average value of three consecutive TEWL readings was obtained from the back skin regions of each mouse.

### Calcium imaging, SOCE measurements, *in vitro* cell scratch assay, and cell proliferation assay

Primary tail keratinocytes were prepared for calcium imaging and SOCE measurements according to a previously published protocol.^45^ (See the Supplemental Methods section for details.) Subsequently, the cells were washed with phosphate-buffered saline (PBS), loaded with 5 µM Fluo4 or Fura2 in Hank’s balanced salt solution (HBSS(+)) at 37°C for 1 hour, and washed with HBSS(−). Fluorescence images were acquired to visualize the fluorescence intensity of Fluo4 using ImageXpress with a FITC filter (Molecular Devices, San Jose, CA). A FlexStation3 (Molecular Devices, San Jose, CA) was used to monitor SOCE by measuring the fluorescence intensity ratio (ratio = F λex 340 nm/F λex 380 nm) of Fura2. After starting the observation or measurement, 2 mM thapsigargin (TG) and 2 mM CaCl_2_ (Fujifilm Wako) were added at 50 seconds and 750 seconds, respectively.

The keratinocytes were subjected to an *in vitro* scratch assay after calcium-induced differentiation. Wound closure was monitored microscopically for up to 12 hours. Cell proliferation was assessed daily using cell counts throughout the 4-day culture period. See the Supplemental Methods section for details.

### *In vivo* wound healing assay

Four-millimeter full-thickness skin wounds were created on the backs of mice using a biopsy punch (BP-40F, KAI Medical, Tokyo, Japan) after hair depilation. To calculate the wound closure rate, photographs of the wounds, including the scale, were acquired every two days throughout the seven-day recovery period.

### Quantitative Polymerase Chain Reaction (qPCR) and RNA-seq

The dorsal whole skin was dissected after shaving, and incubated in 3.8% sodium thiocyanate (Nacalai, Kyoto, Japan) for 20 minutes at room temperature, when necessary to separate the epidermis from the dermis. The separated epidermal layer was washed with PBS and homogenized in a suitable buffer with glass pestles for further experiments. RNA isolation, reverse transcription, and qPCR analysis were performed as described in the Supplementary Materials. All primer sequences are listed in the Supplementally Table 1. RNA-seq analysis was performed by DNAFORM (Yokohama, Japan). Gene ontology (GO) analysis was performed using the Enrichr platform ^46^. See the Supplemental Methods section for details.

### Western blotting

Epidermal protein extracts were prepared and subjected to sodium dodecyl sulfate– polyacrylamide gel electrophoresis (SDS–PAGE) followed by immunoblotting using antibodies against Stim1, Stim2, and α-tubulin. See the Supplemental Methods section for details.

### Protease activity

Epidermal protease activity was measured as previously described with a slight modification^47^. See the Supplemental Methods section for details.

### Statistics

The Mann–Whitney U test was performed for the qPCR (n = 4–8/genotype), the serine protease activities (n = 10/genotype), Delta F340/380, and area under the curve (AUC) for SOCE (n = 6–7/genotype), and TEWL (n = 7–9/genotype). Data represent median values with the interquartile ranges. For the other quantitative experiments, a repeated measures two-way analysis of variance (ANOVA) was performed, and data represent mean ± SD (n = 5–6/genotype). A *p-*value < 0.05 was considered statistically significant.

## Acknowledgement

We appreciate the technical assistance from the Research Support Center, Research Center for Human Disease Modeling, Kyushu University Graduate School of Medical Sciences, partially supported by the Mitsuaki Shiraishi Fund for Basic Medical Research.

We would like to thank the faculty members and technical staff in the Department of Dermatology, Graduate School of Medical Sciences, Kyushu University, for their technical assistance and valuable discussions.

## Author contributions

K.M. and N.H. produced the study concept and design; K.M., N.H., G.T., T.N., and I.T. wrote, reviewed, and revised the paper; K.M., N.H., Z.L., M.N., K.M., F.T., and K.Y. developed the methodology and performed the acquisition, analysis, interpretation, and statistical analysis of data; F.T. provided technical and material support. All authors have read and approved the final paper.

## Availability of Data and Materials

RNA sequencing data in this study have been deposited in the NCBI Gene Expression Omnibus (GEO) under accession number GSE306726.

## References

1 Lee SH, Elias PM, Proksch E, Menon GK, Mao-Quiang M, Feingold KR. Calcium and potassium are important regulators of barrier homeostasis in murine epidermis. J Clin Invest 1992; 89: 530–538.

2 Yuspa SH, Kilkenny AE, Steinert PM, Roop DR. Expression of murine epidermal differentiation markers is tightly regulated by restricted extracellular calcium concentrations in vitro. The Journal of cell biology 1989; 109: 1207–1217.

3 Menon GK, Elias PM. Ultrastructural Localization of Calcium in Psoriatic and Normal Human Epidermis. Archives of Dermatology 1991; 127: 57–63.

4 Pallon J, Malmqvist KG, Werner-Linde Y, Forslind B. Pixe analysis of pathological skin with special reference to psoriasis and atopic dry skin. Cell Mol Biol (Noisy-le-grand) 1996; 42: 111–118.

5 Celli A, Sanchez S, Behne M, Hazlett T, Gratton E, Mauro T. The Epidermal Ca2+ Gradient: Measurement Using the Phasor Representation of Fluorescent Lifetime Imaging. Biophysical Journal 2010; 98: 911–921.

6 Hu Z, Bonifas JM, Beech J, Bench G, Shigihara T, Ogawa H et al. Mutations in ATP2C1, encoding a calcium pump, cause Hailey-Hailey disease. Nat Genet 2000; 24: 61–65.

7 Dhitavat J, Fairclough RJ, Hovnanian A, Burge SM. Calcium pumps and keratinocytes: lessons from Darier’s disease and Hailey–Hailey disease. British Journal of Dermatology 2004; 150: 821–828.

8 Hiramoto K, Kobayashi H, Yamate Y, Ishii M, Sato EF. Intercellular pathway through hyaluronic acid in UVB -induced inflammation. Experimental Dermatology 2012; 21: 911–914.

9 Savignac M, Simon M, Edir A, Guibbal L, Hovnanian A. SERCA2 Dysfunction in Darier Disease Causes Endoplasmic Reticulum Stress and Impaired Cell-to-Cell Adhesion Strength: Rescue by Miglustat. Journal of Investigative Dermatology 2014; 134: 1961–1970.

10 Suryawanshi H, Dhobley A, Sharma A, Kumar P. Darier disease: A rare genodermatosis. Journal of Oral and Maxillofacial Pathology 2017; 21: 321–321.

11 Li F, Adase CA, Zhang L. Isolation and Culture of Primary Mouse Keratinocytes from Neonatal and Adult Mouse Skin. JoVE 2017; : 56027.

12 Park K, Lee SE, Shin K, Uchida Y. Insights into the role of endoplasmic reticulum stress in skin function and associated diseases. The FEBS Journal 2019; 286: 413– 425.

13 Leinonen PT, Myllyla RM, Hagg PM, Tuukkanen J, Koivunen J, Peltonen S et al. Keratinocytes cultured from patients with Hailey-Hailey disease and Darier disease display distinct patterns of calcium regulation. Br J Dermatol 2005; 153: 113–117.

14 Putney JW. Capacitative calcium entry: from concept to molecules. Immunological Reviews 2009; 231: 10–22.

15 Srikanth S, Gwack Y. Orai1, STIM1, and their associating partners. The Journal of Physiology 2012; 590: 4169–4177.

16 Yuan JP, Zeng W, Huang GN, Worley PF, Muallem S. STIM1 heteromultimerizes TRPC channels to determine their function as store-operated channels. Nat Cell Biol 2007; 9: 636–645.

17 Feske S. ORAI1 and STIM1 deficiency in human and mice: roles of store-operated Ca^2+^ entry in the immune system and beyond. Immunological Reviews 2009; 231: 189–209.

18 Putney JW, Steinckwich-Besançon N, Numaga-Tomita T, Davis FM, Desai PN, D’Agostin DM et al. The functions of store-operated calcium channels. Biochimica et Biophysica Acta (BBA) - Molecular Cell Research 2017; 1864: 900–906.

19 Gwack Y, Srikanth S, Oh-hora M, Hogan PG, Lamperti ED, Yamashita M et al. Hair Loss and Defective T- and B-Cell Function in Mice Lacking ORAI1. Molecular and Cellular Biology 2008; 28: 5209–5222.

20 Furukawa Y, Haruyama N, Nikaido M, Nakanishi M, Ryu N, Oh-Hora M et al. Stim1 Regulates Enamel Mineralization and Ameloblast Modulation. J Dent Res 2017; 96: 1422–1429.

21 Lane E. Keratin intermediate filaments and diseases of the skin. In: Madame Curie Bioscience Database [Internet]. Landes Bioscience: Austin (TX), 2000https://www.ncbi.nlm.nih.gov/books/NBK6247/ (accessed 30 July2025).

22 Schmitz A, Lazi’ć E, Koumaki D, Kuonen F, Verykiou S, Rübsam M. Assessing the In Vivo Epidermal Barrier in Mice: Dye Penetration Assays. Journal of Investigative Dermatology 2015; 135: 1–4.

23 Schmitz A, Lazi’ć E, Koumaki D, Kuonen F, Verykiou S, Rübsam M. Assessing the In Vivo Epidermal Barrier in Mice: Dye Penetration Assays. Journal of Investigative Dermatology 2015; 135: 1–4.

24 Komatsu N, Takata M, Otsuki N, Toyama T, Ohka R, Takehara K et al. Expression and Localization of Tissue Kallikrein mRNAs in Human Epidermis and Appendages. Journal of Investigative Dermatology 2003; 121: 542–549.

25 Watt FM. Terminal differentiation of epidermal keratinocytes. Current Opinion in Cell Biology 1989; 1: 1107–1115.

26 Nemes Z, Steinert PM. Bricks and mortar of the epidermal barrier. Exp Mol Med 1999; 31: 5–19.

27 Stanisz H, Mitteldorf C, Bennemann A, Schön MP, Frank J. Subcellular compartmentalization of STIM1 for the distinction of Darier disease from Hailey-Hailey disease. J Deutsche Derma Gesell 2022; 20: 1613–1619.

28 Darbellay B, Arnaudeau S, Ceroni D, Bader CR, Konig S, Bernheim L. Human Muscle Economy Myoblast Differentiation and Excitation-Contraction Coupling Use the Same Molecular Partners, STIM1 and STIM2. Journal of Biological Chemistry 2010; 285: 22437–22447.

29 Liou J, Kim ML, Do Heo W, Jones JT, Myers JW, Ferrell JE et al. STIM Is a Ca2+ Sensor Essential for Ca2+-Store-Depletion-Triggered Ca2+ Influx. Current Biology 2005; 15: 1235–1241.

30 Lu W, Wang J, Peng G, Shimoda LA, Sylvester JT. Knockdown of stromal interaction molecule 1 attenuates store-operated Ca^2+^ entry and Ca^2+^ responses to acute hypoxia in pulmonary arterial smooth muscle. American Journal of Physiology-Lung Cellular and Molecular Physiology 2009; 297: L17–L25.

31 Oh-hora M, Yamashita M, Hogan PG, Sharma S, Lamperti E, Chung W et al. Dual functions for the endoplasmic reticulum calcium sensors STIM1 and STIM2 in T cell activation and tolerance. Nat Immunol 2008; 9: 432–443.

32 Schuhmann MK, Stegner D, Berna-Erro A, Bittner S, Braun A, Kleinschnitz C et al. Stromal Interaction Molecules 1 and 2 Are Key Regulators of Autoreactive T Cell Activation in Murine Autoimmune Central Nervous System Inflammation. The Journal of Immunology 2010; 184: 1536–1542.

33 Numaga-Tomita T, Putney JW. Role of STIM1- and Orai1-mediated Ca2+ entry in Ca2+-induced epidermal keratinocyte differentiation. Journal of Cell Science 2013; 126: 605–612.

34 Vandenberghe M, Raphaël M, Lehen’kyi V, Gordienko D, Hastie R, Oddos T et al. ORAI1 calcium channel orchestrates skin homeostasis. Proc Natl Acad Sci USA 2013; 110. doi:10.1073/pnas.1310394110.

35 Raiko L, Leinonen P, Hägg P, Peltonen J, Oikarinen A, Peltonen S. Tight junctions in Hailey-Hailey and Darier’s diseases. Dermatol Reports 2009; 1: e1.

36 Raiko L, Siljamäki E, Mahoney MG, Putaala H, Suominen E, Peltonen J et al. Hailey–Hailey disease and tight junctions: Claudins 1 and 4 are regulated by ATP 2C1 gene encoding Ca^2+^ /Mn^2+^ ATP ase SPCA 1 in cultured keratinocytes. Experimental Dermatology 2012; 21: 586–591.

37 Li N, Park M, Xiao S, Liu Z, Diaz LA. ER -to-Golgi blockade of nascent desmosomal cadherins in SERCA2 -inhibited keratinocytes: Implications for Darier’s disease. Traffic 2017; 18: 232–241.

38 Brattsand M, Egelrud T. Purification, Molecular Cloning, and Expression of a Human Stratum Corneum Trypsin-like Serine Protease with Possible Function in Desquamation. Journal of Biological Chemistry 1999; 274: 30033–30040.

39 Kishibe M, Bando Y, Terayama R, Namikawa K, Takahashi H, Hashimoto Y et al. Kallikrein 8 Is Involved in Skin Desquamation in Cooperation with Other Kallikreins. Journal of Biological Chemistry 2007; 282: 5834–5841.

40 Caubet C, Jonca N, Brattsand M, Guerrin M, Bernard D, Schmidt R et al. Degradation of Corneodesmosome Proteins by Two Serine Proteases of the Kallikrein Family, SCTE/KLK5/hK5 and SCCE/KLK7/hK7. Journal of Investigative Dermatology 2004; 122: 1235–1244.

41 Billi AC, Ludwig JE, Fritz Y, Rozic R, Swindell WR, Tsoi LC et al. KLK6 expression in skin induces PAR1-mediated psoriasiform dermatitis and inflammatory joint disease. Journal of Clinical Investigation 2020; 130: 3151–3157.

42 Morizane S, Yamasaki K, Kabigting FD, Gallo RL. Kallikrein Expression and Cathelicidin Processing Are Independently Controlled in Keratinocytes by Calcium, Vitamin D3, and Retinoic Acid. Journal of Investigative Dermatology 2010; 130: 1297–1306.

43 Kobashi M, Morizane S, Sugimoto S, Sugihara S, Iwatsuki K. Expression of serine protease inhibitors in epidermal keratinocytes is increased by calcium but not 1,25-dihydroxyvitamin D_3_ or retinoic acid. Br J Dermatol 2017; 176: 1525–1532.

44 Tominaga M, Ozawa S, Tengara S, Ogawa H, Takamori K. Intraepidermal nerve fibers increase in dry skin of acetone-treated mice. Journal of Dermatological Science 2007; 48: 103–111.

45 Li F, Adase CA, Zhang L. Isolation and Culture of Primary Mouse Keratinocytes from Neonatal and Adult Mouse Skin. .

46 Xie Z, Bailey A, Kuleshov MV, Clarke DJB, Evangelista JE, Jenkins SL et al. Gene Set Knowledge Discovery with Enrichr. Current Protocols 2021; 1: e90.

47 Kishibe M, Bando Y, Terayama R, Namikawa K, Takahashi H, Hashimoto Y et al. Kallikrein 8 Is Involved in Skin Desquamation in Cooperation with Other Kallikreins. Journal of Biological Chemistry 2007; 282: 5834–5841.

48 Furuse M, Hata M, Furuse K, Yoshida Y, Haratake A, Sugitani Y et al. Claudin-based tight junctions are crucial for the mammalian epidermal barrier. The Journal of Cell Biology 2002; 156: 1099–1111.

