## Supplemental Methods for "Impaired store-operated Ca^2+^ entry in mouse epidermis leads to reduced epidermal barrier function via Klk activation"

**Immunohistochemistry**

Immunohistochemistry for Stim1 (1:50, HPA012123, Merck, Darmstadt, Germany), Dsg1 (Desmograin1) (1:500, 24587-1-AP, Proteintech, Rosemont, IL), Cdh1 (E-cadherin) (1:500, #3195, Cell signaling, Danvers, MA), Flg (filaggrin) (1:100, ab234406, Abcam, Waltham, MA), and Ki67 (1:1000, 28074-1-AP, Proteintech) was conducted after antigen retrievals in citrate buffer and blocking with 5% sheep serum for 15 minutes. Samples were incubated with a secondary antibody (anti-rabbit IgG antibody with Alexa Fluor 488, ab150077, Abcam) and mounted with DAPI-containing mounting medium (Prolong Gold with DAPI, Thermo Fisher Scientific, Waltham, MA).

**Lucifer yellow assay**

Mice (6-to-7-week-old) were restrained in Petri dishes with their backs in contact with 1 mM Lucifer yellow (Wako) in Ringer’s solution (pH 7.4) at 37°C. (PNAS 1998 95:1044-1049). After 1 hour of incubation, the dorsal skin was excised and immediately frozen on dry ice. The sections for Lucifer yellow were fixed in 4% PFA for 10 minutes before mounting with DAPI-containing medium.

***Lac Z* staining**

The sections from post-natal day 7 neonatal mice were fixed in 4% PFA for 10 minutes, and *LacZ* activity was visualized using a β-gal assay kit (K1465-01, Thermo Fisher Scientific), followed by counterstaining with eosin.

**Biotin permeability assay**

The biotin labeling assay was performed with a protocol adapted from Furuse et al.^48^ Briefly, 50 µl of 10 mg/ml Sulfo-NHS-LC-Biotin (B1022, Merck) was dissolved in PBS containing 1 mM CaCl₂, which was then injected subcutaneously. After a 30-minute incubation period, the dorsal skin was excised and immediately frozen on dry ice. The frozen sections were fixed in 4% PFA for 10 minutes, followed by fixation in 95% ethanol for 20 minutes at 4°C. After blocking with 5% sheep serum for 15 minutes, the sections were incubated with Texas Red-conjugated streptavidin (1:200, SA-5006, Vector Laboratories, Newark, CA) for 1 hour and mounted with DAPI-containing medium. All sections were observed with a fluorescence microscope (BZ-X700, Keyence, Tokyo, Japan).

**Calcium imaging and SOCE measurements using primary tail keratinocytes**

After the tail skin was serially digested with dispase and trypsin (Thermo Fisher Scientific), the low calcium keratinocyte growth medium (# 06-174, Lonza, Basel, Switzerland) supplemented with 8% (vol/vol) chelexed serum was added to the isolated epidermis to produce a suspension of individual cells. The isolated epidermal keratinocytes were seeded at 2 × 10^4^ cells/well on collagen I-coated 96-well plates (356649, Corning, NY) and incubated at 37°C in a humidified incubator with 5% CO_2_. Calcium imaging of the keratinocytes was performed using intracellular Ca²⁺ measurement kits (Calcium Kit–Fura2 or Fluo4, DOJINDO, Kumamoto, Japan) according to the manufacturer's protocol.

***In vitro* cell scratch assay and cell proliferation assay**

Primary keratinocytes isolated from mouse tails mentioned above were seeded in 6-well plates at 10 × 10^6^ cells per well and cultured until confluent for the *in vitro* cell scratch assay. The culture medium was switched from a low calcium keratinocyte growth medium to a high calcium differentiation medium containing 1.8 mM CaCl_2_ before scratching. A 200 μL pipette tip was used to scrape across the dish, and the resulting wound was washed with PBS. Phased images were captured using a microscope (OLYMPUS iX71) every 3 hours for 12 hours after wounding. The wound and scratch area were quantified using ImageJ software (NIH, Bethesda, MD). To assess cell proliferation, the keratinocytes were seeded in 24-well plates at 3 × 10⁴ cells per well and cultured in a low calcium keratinocyte growth medium supplemented with 0.03 mM Ca²⁺ for four days. The cell numbers per well were directly counted daily with an automated cell counter (LUNA-II, Logos Biosystems, South Korea).

**qPCR**

Total RNA was extracted using TRIzol reagent (Thermo Fisher Scientific) and purified with the RNeasy Mini Kit (Qiagen, Valencia, CA). Reverse transcription was performed according to the manufacturer's protocol for cDNA synthesis (ReverTra Ace, TOYOBO, Osaka, Japan) for qPCR. The qPCR procedure was performed using a qPCR kit (KAPA SYBR Fast, Kapa Biosystems, Wilmington, MA) coupled to a qPCR detection system (CFX Connect, Bio-Rad, Hercules, CA). The expression levels were evaluated using the ΔΔCt method, first normalized to the housekeeping gene *Hprt* and then to the *Stim1/2^fl/fl^* mouse as a control (n = 4–8).

**RNA-seq**

After assessing the RNA quality (Bioanalyzer2100, Agilent)*,* RNA-seq libraries were prepared using the QIAseq UPX 3' Transcriptome Kit (#333088, QIAGEN). Subsequently, the libraries were adapted for the MGI sequencing platform with the MGIEasy Universal Library Conversion Kit (App-A) (#1000004155, MGI) according to the manufacturer's protocols. The libraries were sequenced using a DNBSEQ-G400 instrument (MGI). Subsequently, the obtained reads were mapped to the mouse GRCm39 genome using STAR (version 2.7.10a). Reads on annotated genes were counted using featureCounts (version 2.0.1). Clustering analysis was performed using the Subio platform (Subio, Nagoya, Japan). Genes more than two-fold differentially expressed between the groups were used for analysis.

**Western blotting**

The proteins from the separated epidermal layer were extracted in 8 M urea (Fujifilm Wako) and centrifuged at 15,000 g for 20 minutes. The supernatant was collected and stored at −80°C until use. A total mass of 15 µg of protein from the epidermal skin extract was separated on 4–12% Bis-Tris gels in the MOPS running buffer (NuPage, Thermo Fisher Scientific). The separated proteins were transferred to nitrocellulose membranes and blocked in PBS containing 0.1% Tween-20 and 5% skim milk for 1 hour at room temperature. The blocked membranes were incubated overnight at 4°C with antibodies for Stim1 (1:1000, HPA012123), Stim2 (1:1000, ACC-064, Alomone, Jerusalem, Israel), or α-tubulin (1:3000, T9026, Merck) in the blocking solution. After incubation with goat anti-rabbit or goat anti-mouse IgG-HRP (1:1000, SC-2004, or SC-2005; Santa Cruz Biotechnology) as secondary antibodies for 1 hour, proteins were visualized and recorded using an image reader (LAS-3000, Fujifilm, Tokyo, Japan).

**Protease activity**

Shaved dorsal skin was dissected and incubated in 1.8 M NaCl for 3 hours at 37°C, and the epidermis was separated at the dermal–epidermal junction. The epidermis was homogenized in extraction buffer containing 60 mM Tris-HCl (pH 8.0), 150 mM NaCl, 1% Nonidet P-40, 5 mM EDTA in a glass pestle and centrifuged at 15,000g for 20 minutes at 4°C. A total mass of 5 µg protein from the epidermis was adjusted to a total volume of 200 µl with 100 µM synthetic substrates for trypsin (Boc-Phe-Ser-Arg-MCA, 3107-V, Peptide Institute, Osaka, Japan) or chymotrypsin (Suc-Leu-Leu-Val-Tyr-MC, 3120-V, Peptide Institute) and added to a black 96-well plate (23302, Berthold Technologies, Bad Wildbad, Germany). As a negative control, 1 µM PMSF (#8553, Cell Signaling) was added to the samples. Fluorescence was measured with the FlexStation3 at wavelengths (λex=380 nm, λem=460 nm) after 5 hours of incubation. The fluorescence intensity of the negative control was subtracted from the samples without PMSF and normalized to the *Stim1/2^fl/fl^* mouse.
