## Supplemental Table for "Impaired store-operated Ca^2+^ entry in mouse epidermis leads to reduced epidermal barrier function via Klk activation"

**Supplemental Table 1**

qPCR primers

| ***Primer*** | ***Sequences*** | ***GenBank accession*** |
| --- | --- | --- |
| ***Stim1 (F)***  ***Stim1 (R)***  ***Stim2 (F)***  ***Stim2 (R)***  ***Orai1 (F)***  ***Orai1 (R)***  ***Dsg1a (F)***  ***Dsg1a (R)***  ***Chd1 (F)***  ***Chd1(R)***  ***Flg (F)***  ***Flg (R)***  ***Klk5 (F)***  ***Klk5 (R)***  ***Klk6 (F)***  ***Klk6 (R)***  ***Klk7 (F)***  ***Klk7 (R)***  ***Spink5 (F)***  ***Spink5 (R)***  ***Hprt (F)***  ***Hprt (R)*** | 5’-TGAAGAGTCTACCGAAGCAGA-3’  5’-AGGTGCTATGTTTCACTGTTGG-3’  5’-CGAAGTGGACGAGAGTGATGA-3’  5’-GGAGTGTTGTTCCCTTCACATT-3’  5’-GATCGGCCAGAGTTACTCCG-3’  5’-TGGGTAGTCATGGTCTGTGTC-3’  5’-CAAGGCACTTCTTCCACTGAGA-3’  5’-CGCTGCCTCCCCATGA-3’  5’-ATCAGCTGCCCCGAAAATGA-3’  5’-ACTTTCAGCCAGCCTGTCTC-3’  5’-AGACTGGGAGGCAAGCTACA-3’  5’-CCTGCCTCCTTCAGAGTCAC-3’  5’-ATGGGCAATGGCTACCCTG-3’  5’-GTTCGGTTCCAGAGGGGTT-3’  5’-GCCCTCTACACCTCAGGTCA-3’  5’-ATCACCTGCAGATTCGGTTT-3’  5’-CGTCATGTACCGTCTCTGGA-3’  5’-CACTCCCTGGAGGAGATGAG-3’  5’-CACTGACCCAGCAAAGTTGA-3’  5’-TTTTCGCATTCTCAGCACAC-3’  5’-TCAGTCAACGGGGGACATAAA-3’  5’-GGGGCTGTACTGCTTAACCAG-3’ | NM_009287  NM_001081103  NM_175423  [NM_010079](https://www.ncbi.nlm.nih.gov/entrez/viewer.fcgi?db=nucleotide&id=2649350490)  NM_009864  XM_017319842.1  NM_026806  NM_011177  NM_011872  NM_001081180  NM_013556 |

**Note:** These primer sequences were obtained from the following manuscripts: *Stim1, Stim2, Orai1, Hprt* [1], *Dsg1a* [2], *Klk5* [3]*, Klk6* [4], *Spink5* [5]. The primers for *Chd1, Flg*, and *Klk7* were designed through the online tool Primer-BLAST (https://www.ncbi.nlm.nih.gov/tools/primer-blast/).

[1] Furukawa Y, Haruyama N, Nikaido M, Nakanishi M, Ryu N, Oh-Hora M, Kuremoto K, Yoshizaki K, Takano Y, Takahashi I. Stim1 Regulates Enamel Mineralization and Ameloblast Modulation. J Dent Res. 2017 Nov;96(12):1422-1429. doi: 10.1177/0022034517719872. Epub 2017 Jul 21. PMID: 28732182.

[2] Godsel LM, Roth-Carter QR, Koetsier JL, Tsoi LC, Huffine AL, Broussard JA, Fitz GN, Lloyd SM, Kweon J, Burks HE, Hegazy M, Amagai S, Harms PW, Xing X, Kirma J, Johnson JL, Urciuoli G, Doglio LT, Swindell WR, Awatramani R, Sprecher E, Bao X, Cohen-Barak E, Missero C, Gudjonsson JE, Green KJ. Translational implications of Th17-skewed inflammation due to genetic deficiency of a cadherin stress sensor. J Clin Invest. 2022 Feb 1;132(3):e144363. doi: 10.1172/JCI144363. PMID: 34905516; PMCID: PMC8803337.

[3] Kidana K, Tatebe T, Ito K, Hara N, Kakita A, Saito T, Takatori S, Ouchi Y, Ikeuchi T, Makino M, Saido TC, Akishita M, Iwatsubo T, Hori Y, Tomita T. Loss of kallikrein-related peptidase 7 exacerbates amyloid pathology in Alzheimer’s disease model mice. EMBO Mol Med. 2018 Mar;10(3):e8184. doi: 10.15252/emmm.201708184. PMID: 29311134; PMCID: PMC5840542.

[4] Hwang YS, Cho HJ, Park ES, Lim J, Yoon HR, Kim JT, Yoon SR, Jung H, Choe YK, Kim YH, Lee CH, Kwon YT, Kim BY, Lee HG. KLK6/PAR1 Axis Promotes Tumor Growth and Metastasis by Regulating Cross-Talk between Tumor Cells and Macrophages. Cells. 2022 Dec 16;11(24):4101. doi: 10.3390/cells11244101. PMID: 36552865; PMCID: PMC9777288.

[5] Patel S, Xi ZF, Seo EY, McGaughey D, Segre JA. Klf4 and corticosteroids activate an overlapping set of transcriptional targets to accelerate in utero epidermal barrier acquisition. Proc Natl Acad Sci U S A. 2006 Dec 5;103(49):18668-73. doi: 10.1073/pnas.0608658103. Epub 2006 Nov 27. PMID: 17130451; PMCID: PMC1693720.

**Supplemental Table 2**

**GO biological process**

| **GO Term** | ***p-*value** | **Odds Ratio** | **related genes** |
| --- | --- | --- | --- |
| Intermediate Filament Organization (GO:0045109) | 1.68E-08 | 44.09534368 | Krt17;Tchh;Krt79;Krt75;Krt6b;Krt6a |
| Supramolecular Fiber Organization (GO:0097435) | 1.36E-04 | 8.622280817 | Krt17;Tchh;Krt79;Krt75;Krt6b;Krt6a |
| Protein Autoprocessing (GO:0016540) | 7.19E-04 | 59.07555556 | Ctsl;Klk6 |
| Epidermis Development (GO:0008544) | 0.001085724 | 16.32256298 | Fabp5;Col7a1;Klk6 |
| Lipid Transport (GO:0006869) | 0.002376366 | 12.29938017 | Fabp5;Npc2;Bltp1 |
| Collagen Catabolic Process (GO:0030574) | 0.002406549 | 30.5348659 | Ctsl;Klk6 |
| Phospholipid Transport (GO:0015914) | 0.009056603 | 14.98606403 | Slc66a2;Npc2 |
| Skin Development (GO:0043588) | 0.011472991 | 13.19137645 | Flg;Dhcr24 |
