## Supplemental Material for "Impaired store-operated Ca^2+^ entry in mouse epidermis leads to reduced epidermal barrier function via Klk activation"

Fig. 1d

Stim1

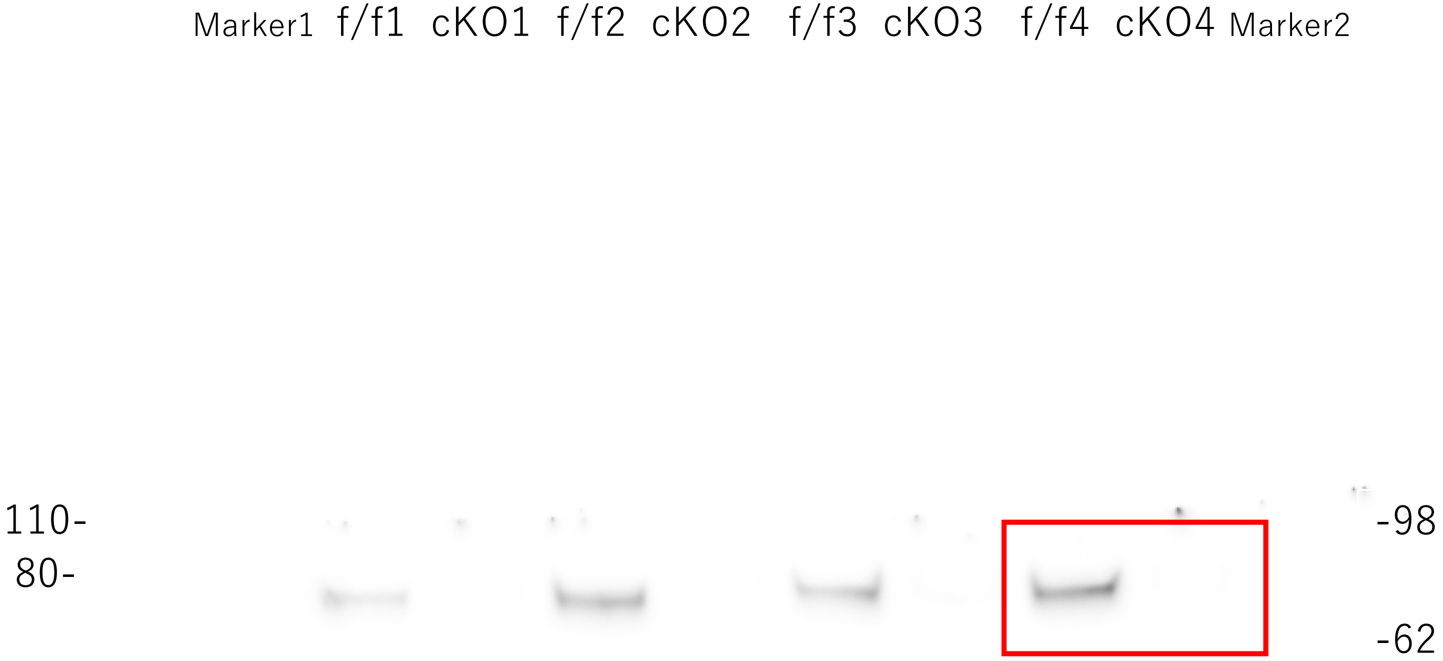


The lanes f/f4 and cKO4 were used in the Fig. 1d.

The images was digitized using BioTools MultiImager II ChemiBOX.

Marker1: Sharp Pre-stained Protein Standard, Marker2: Seeblue Pre-stained Protein Standard.

Supplemental Material

Fig. 1d

Stim2

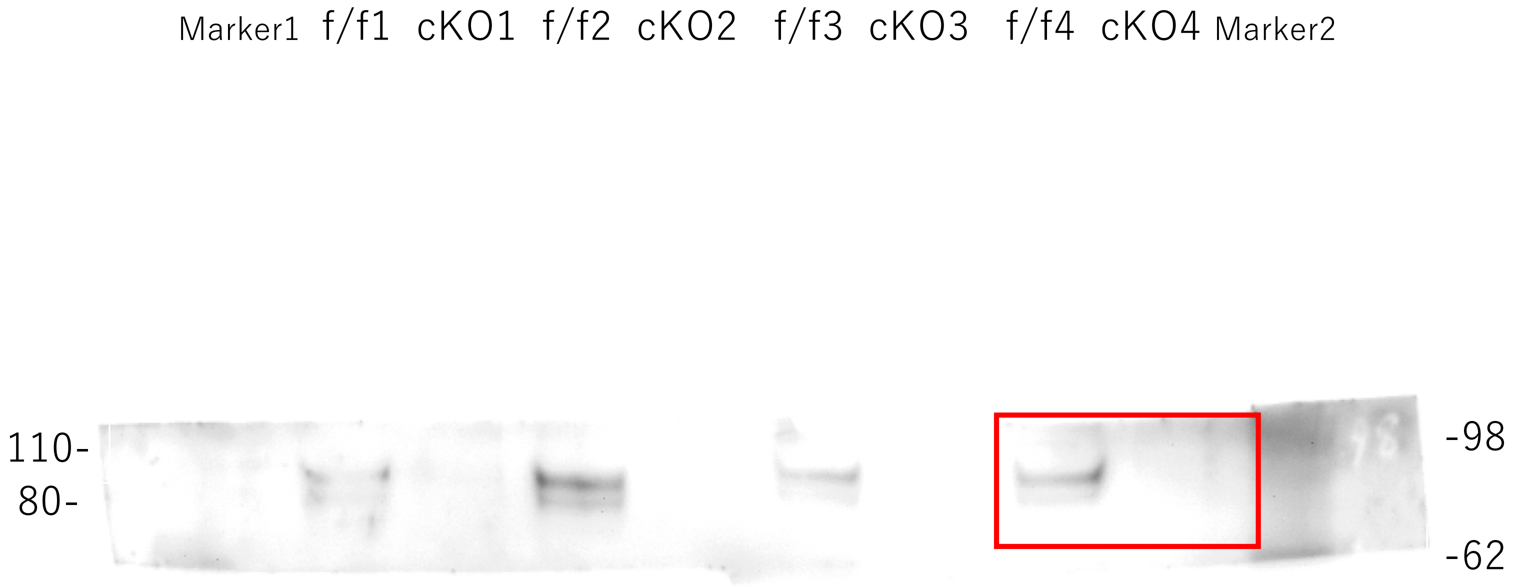


The lanes f/f4 and cKO4 were used in the Fig. 1d.

The images was digitized using BioTools MultiImager II ChemiBOX.

Marker1: Sharp Pre-stained Protein Standard, Marker2: Seeblue Pre-stained Protein Standard.

Supplemental Material

Fig. 1d

α-tubulin

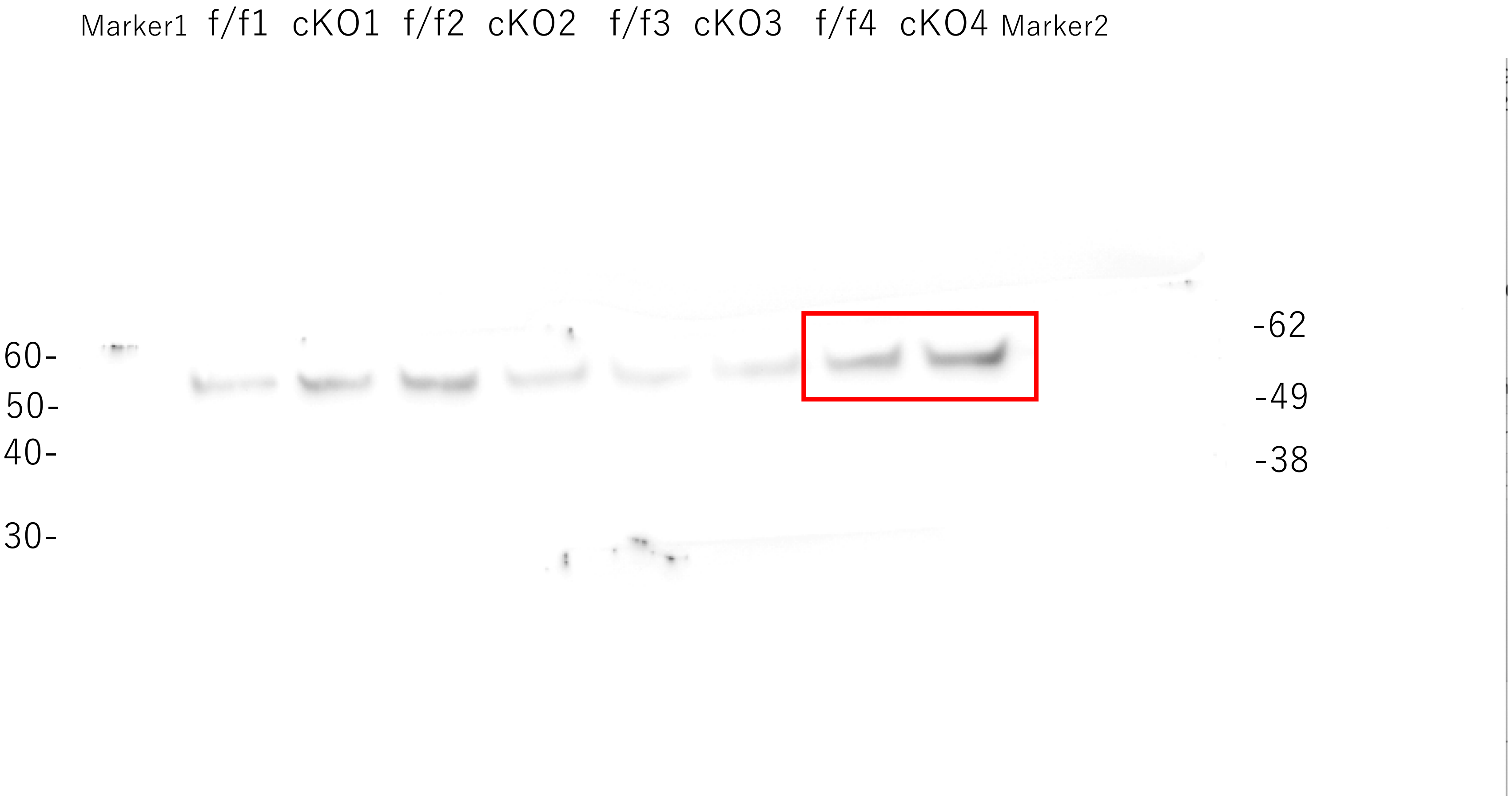


The lanes f/f4 and cKO4 were used in the Fig. 1d.

The images was digitized using BioTools MultiImager II ChemiBOX.

Marker1: Sharp Pre-stained Protein Standard, Marker2: Seeblue Pre-stained Protein Standard.
